# SHANK3 downregulation in the VTA accelerates the extinction of contextual associations induced by non-familiar conspecific interaction

**DOI:** 10.1101/342295

**Authors:** Sebastiano Bariselli, Alessandro Contestabile, Stamatina Tzanoulinou, Camilla Bellone

## Abstract

Conditioned place preference (CPP) paradigms, traditionally adopted to study the reinforcing properties of drugs of abuse, have been developed to investigate the neurobiological mechanisms underlying the reinforcing properties of social stimuli. These protocols are largely based on single-housing before and/or during the social stimulus-contextual cues acquisition phase. Here, based on previously established social interaction-induced CPP paradigms, we characterize a place preference task relying on the reinforcing properties of free interaction with a non-familiar (novel) conspecific. The formation of contextual associations induced by the interaction with a novel social stimulus does not require single-housing, necessitates dopamine (DA) receptor 2/3 (D2/3R) activation and undergoes extinction. Interestingly, while extinction of CPP responses is reduced by single-housing, it is accelerated by the downregulation of the autism spectrum disorder (ASD)-related protein *SHANK3* in the ventral tegmental area (VTA). Thus, inspired by the literature on drug of abuse-induced contextual learning, we propose that acquisition and extinction of CPP might be used as behavioral assays to assess social-induced contextual association and “social-seeking” dysfunctions in animal models of psychiatric disorders.

## 1 Introduction

Over the years, several experimental paradigms have been developed to define the different aspects of social behaviour (Sandi and Haller, 2015), such as sociability (Nadler et al., 2004), preference for social novelty (Nadler et al., 2004), social memory and novelty recognition (Kogan et al., 2000; Ferguson et al., 2000). During dyadic encounters, social interactions might acquire positive valence and promote approach behavior (Kogan et al., 2000; Ferguson et al., 2000) or negative valence and drive avoidance behavior (Golden et al., 2011). Moreover, rodents can learn to associate certain environmental cues (conditioned stimuli, CS) with the availability of positive social interaction (unconditioned stimuli, US) resulting in conditioned place preference (CPP) responses. CPP tasks, classically used to study the reinforcing properties of drugs of abuse (Tzschentke, 2007), have been adopted to test the ability of conspecific interaction to promote contextual associative learning. In fact, different animal species, such as rats (Thiel et al., 2008; Trezza et al., 2009; Calcagnetti and Schechter, 1992; Crowder and Hutto, 1992; Van den Berg et al., 1999; Kummer et al., 2011), mice (Panksepp and Lahvis, 2007; Dölen et al., 2013; Hung et al., 2017) and squirrels (Lahvis et al., 2015), show social-induced CPP responses. Although the number or the duration of conditioning sessions (Trezza et al., 2009; Thiel et al., 2008), the genetic background (Panksepp et al., 2007; Pinheiro et al., 2016) and the age of the animals (Douglas 2004) influence place preference acquisition, one common aspect among all the different social-induced CPP protocols is that the animals are single-housed before and/or during conditioning (Calcagnetti and Schechter, 1992; Kummer et al., 2011; Fritz et al., 2011; Pinheiro et al., 2016; Molas et al., 2017; Thiel et al., 2008). Single-housing, possibly through adaptations occurring within the dopaminergic system (Lewis et al., 1994; Fabricius et al., 2011; Whitaker et al., 2013; Lallai et al., 2016), enhances sociability (Panksepp and Beatty, 1980; Niesink and Van Ree, 1989; Vanderschuren et al., 1995), increases the incentive value of social stimuli (Van den Berg et al., 1999; Trezza et al., 2009) and seems necessary to obtain social-induced CPP (Douglas et al., 2004; Trezza et al., 2009).

The reward system originates from the highly heterogeneous dopamine (DA) neurons of the ventral tegmental area (VTA) that project to several brain regions, such as the nucleus accumbens (NAc) and the prefrontal cortex (Lammel et al., 2014; Bariselli et al., 2016a; Morales and Margolis, 2017). Despite their diversity, at least a subclass of midbrain DA neurons increase their activity in response to unexpected rewards and, after associative learning, to cues predicting reward availability (Schultz et al., 1997; Tobler et al., 2005; Cohen et al., 2012). In respect to social behaviour, VTA DA neuron activation is both necessary and sufficient to elicit interaction among peers (Gunaydin et al., 2014) and the function of DA receptor 1 and 2 (D1R and D2R) is fundamental to express appropriate social behaviour in rodents (Gunaydin et al., 2014, Lee et al., 2017). The reward system also plays a fundamental role in associative learning. Indeed, while optogenetic activation of DA neurons promotes CPP and reinforces instrumental learning (Tsai et al., 2009; Adamantidis et al., 2011), alterations of D2R/D3R function perturbs the acquisition of reward-induced place preference (Vidal-Infer et al., 2012; Hoffman and Beninger, 1989; Rezayof et al., 2002; Le Foll et al., 2005; Cunningham et al., 2000). Emerging evidence suggests the involvement of the DA system in social-induced learning of new contextual associations. In fact, social-induced CPP activates several regions within the reward system (Rawas et al., 2012), it is controlled by oxytocin within the VTA and NAc (Hung et al., 2017; Dölen et al., 2013) and is impaired by the DA re-uptake inhibitor, methylphenidate (Trezza et al., 2009).

Associative learning is subject to extinction, a behavioral phenomenon caused by the repeated presentations of contextual cues in the absence of the reinforcer and characterized by a timely reduction of the conditioned responses. The extinction of conditioned responses could depend on the formation of a new memory, by which contextual cues previously associated with a reinforcer become associated with its absence (for a comprehensive view on extinction theories see Dunsmoor et al., 2015; Bouton, 2004; Quirk and Mueller, 2008). Interestingly, appropriate synaptic transmission and plasticity at synaptic inputs onto VTA DA neurons is fundamental, not only for the acquisition (Sartor and Aston-Jones, 2012; Mills et al., 2017; Huang et al., 2016; Stuber et al., 2008) but also for the extinction of reinforcement learning (Engblom et al., 2008; Chen and Chen, 2015). However, although it has been shown that social-induced CPP undergoes extinction in adolescent rats (Trezza et al., 2009), the involvement of the VTA in this process remains elusive.

In humans, emerging evidence suggests an existing association between reward system alterations and autism spectrum disorder (ASD; Pavăl, 2017), a neurodevelopmental disorder characterized by social dysfunctions and compulsivity symptoms (*DMS-V*). For example, polymorphisms of the *D2R* and *D3R* genes (Staal et al., 2012; Comings et al., 1991) and increased D2R binding (Nakamura et al., 2010) are reported in ASD cases. Furthermore, in genetic models of ASD, VTA dysfunctions lead to sociability deficits (Bariselli et al., 2016b; Krishnan et al., 2017), suggesting that impairments in the reward system might underlie aberrant social-induced learning. Several genetic mouse lines with *Shank3* gene mutations have been developed to investigate the molecular and neuronal impairments underlying ASD-related alterations in rodents (Bariselli and Bellone, 2016). In addition to impairments in the social and compulsivity domains, *Shank3* KO mutant mice display deficits in associative learning with stimuli of both negative (fear conditioning, Drapeau et al., 2014) and positive (food consumption, Wang et al., 2016) valence. Previously, we have shown that the early post-natal downregulation of *Shank3* in the VTA induces synaptic excitatory deficits associated to social preference (Bariselli et al., 2016b), pointing at an impaired social reward processing. Thus, since VTA neuronal plasticity is necessary for appropriate learning and extinction, we aimed to determine whether *Shank3* deficiency in the VTA would alter both components of social-induced CPP.

In this study, we propose and characterize a behavioural paradigm of social-induced CPP to assess the reinforcing properties of direct social interaction with a non-familiar (novel) conspecific (non-familiar conspecific-induced CPP). We demonstrate that non-familiar conspecific promotes place preference in both single-housed and group-housed C57Bl6/j late adolescence male mice and its acquisition depends on D2/D3R activation. Furthermore, as this form of CPP is expressed even when group-housed mice are conditioned with non-familiar *vs* familiar conspecifics, we conclude that the direct interaction with a non-familiar conspecific is *per se* reinforcing. In addition, while preference can be lost, the retention of the conditioned behavioural responses are heightened by single-housing. Lastly, downregulation of *Shank3* expression in the VTA accelerates extinction without inducing major abnormalities in the acquisition of non-familiar conspecific-induced CPP.

## 2 Materials and methods

### 2.1 Animals

The experimental procedures described here were conducted in accordance with the Swiss laws and previously approved by the Geneva Cantonal Veterinary Authority. Male C57Bl/6j mice were purchased from Charles River Laboratories and housed in the institutional animal facility under standard 12h/12h light/dark cycles with food and water *ad libitum.* Experimental animals were group-housed (2-3 per cage), or single-housed only for the single-housing experimental condition, and tested during late adolescence, at the 8th or 9th week of life. Younger non-familiar mice (3-4 weeks) were single-housed and used as stimuli animals during the conditioning sessions of the CPP protocol.

### 2.2 Conditioned place preference induced by non-familiar conspecific interaction

The apparatus used (Bioseb, Model BIOSEB, In Vivo Research Instruments, spatial place preference box for mice LE895) for the CPP protocol has two square-shaped chambers with different patterns on the walls and different textured floors. The two chambers are interconnected by a small corridor, with transparent walls and floor, and removable doors allow the corridor to be closed (**Figure 1**).

**Figure 1.**
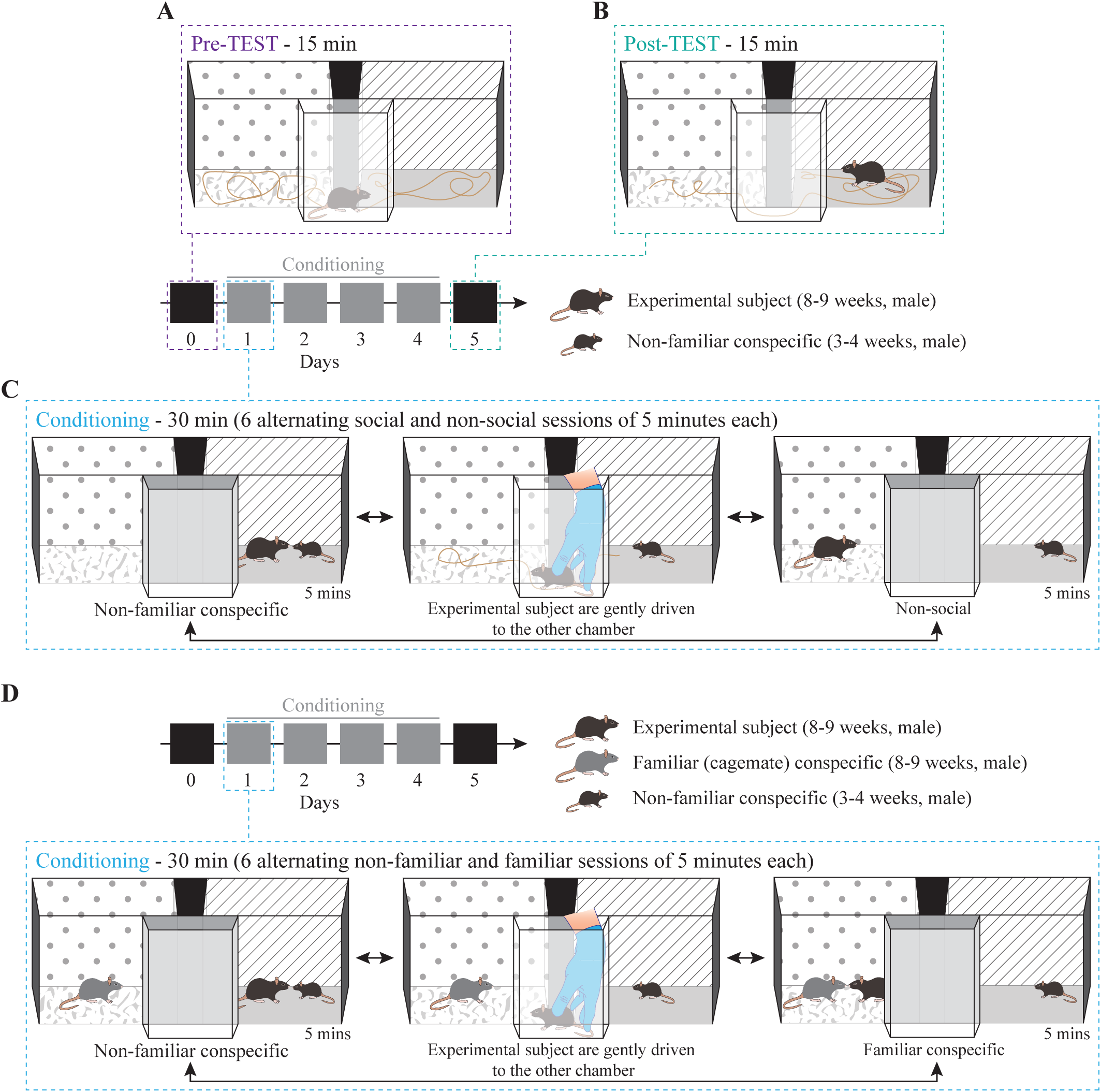
Non-familiar conspecific-induced conditioned place preference: a variation of social-induced CPP to assess the reinforcing properties of social interaction. Schematic 3D representation of the CPP procedure and apparatus. The protocol consisted in: (**A**) 15 minutes pre-TEST, (**B**) 15 minutes post-TEST and (**C**) conditioning phase (30 minutes per day for 4 consecutive days). (**D**) Schematic 3D representation of conditioning phase for non-familiar vs familiar social stimuli protocol, during which the experimental mouse interacted with a non-familiar social stimulus in one chamber and with a familiar (cage-mate) stimulus in the other one.

To note, after each pre-TEST, conditioning session, post-TEST and extinction protocol the arena was cleaned thoroughly with 1% acetic acid and allowed to dry before continuing with the next animal.

*Day 0 - Pre-TEST:* experimental mice freely explored the CPP apparatus for 15 minutes to establish baseline preference for both chambers (**Figure 1A**). On the same day, stimuli mice were habituated either to one or the other chamber for 15 minutes. The chamber that they were habituated in was the chamber that they were in during the conditioning sessions.

*Days 1-4* – *Conditioning, paired protocol (non-familiar vs non-social session):* for each experimental mouse one chamber was randomly assigned as the paired session chamber, with the presence of a non-familiar mouse (US +) and the other as the non-social session chamber (US -). This protocol remained stable for each mouse throughout the conditioning sessions of Days 1-4. Experimental mice were subjected to six alternating conditioning sessions of 5 minutes each, resulting in 30 minutes of total conditioning time per day (e.g. non-familiar mouse session 1, non-social session 1, non-familiar mouse session 2, non-social session 2, non-familiar mouse session 3, non-social session 3) (**Figure 1C**). To counteract bias formation, each day the starting session was counterbalanced across mice and alternated for each mouse, such that, if for example on day 1 the experimental subject was exposed to the paired session first, on day 2 the conditioning started with the non-social session. After the end of each 5-minute trial, the animals were gently guided by the experimenter through the corridor and towards the other chamber (**Figure 1C**), so the animals were exposed to the spatial orientation of both chambers and apparatus.

*Days 1-4* – *Conditioning, contingency break protocol:* this experimental protocol did not maintain the stable contingencies between chamber and stimulus. Specifically, the assignment of one context to the non-familiar stimulus (US +) and another context to the non-social (US -) was inverted each day (e.g. day 1: non-familiar mouse in chamber A vs non-social in chamber B; day 2: non-familiar mouse in chamber B *vs* non-social in chamber A; day 3: non-familiar mouse in chamber A *vs* non-social in chamber B; day 4: non-familiar mouse in chamber B *vs* non-social in chamber A).

*Days 1-4* – *Conditioning, paired protocol (familiar session vs non-familiar session):* for each experimental mouse one chamber was assigned as the paired session chamber, with the presence of the non-familiar mouse (US +), and the other chamber was associated with the presence of a cage-mate, the familiar stimulus (US -). This protocol remained stable for each mouse throughout the conditioning sessions of Days 1-4. Experimental mice were subjected to six alternating conditioning sessions of five minutes each, resulting in 30 minutes of total conditioning time per day (e.g. non-familiar mouse session 1, familiar mouse session 1, non-familiar mouse session 2, familiar mouse session 2, non-familiar mouse session 3, familiar mouse session 3) (**Figure 1D**). Each day the starting session was counterbalanced across mice and alternated for each mouse. After the end of each 5-minute trial, the animals were gently guided by the experimenter through the corridor and towards the other chamber (**Figure 1D**).

We should note that, in this paradigm, although the stimulus mouse represented a stranger stimulus at the first conditioning session, the non-familiarity aspect was maintained since animals returned to their respective home-cages and kept separated for 24 hours. Thus, in this case, the US exposure represented a valued condition. In contrast, when the experimental subject had access to the familiar stimulus both in the home-cage and in the conditioned side of the apparatus, the familiar stimulus mouse exposure represented a devalued condition.

*Day 5- Post-TEST:* twenty-four hours after the last conditioning session, experimental mice were placed in the corridor of the CPP apparatus and, after lifting the removable doors, the animals could freely explore the arena once again for 15 minutes and establish a preference (**Figure 1B**) in the absence of the US.

*Extinction:* starting forty-eight hours after the post-TEST session, experimental mice were exposed for several days (5-6 days) to the empty apparatus. In particular, mice were placed in the corridor of the CPP apparatus and, after lifting the removable doors, explored the arena for 15-minute-long sessions in the absence of the US. After each extinction session, the arena was cleaned thoroughly with 1% acetic acid and allowed to dry before continuing with the next animal.

The behavior of the animals was tracked automatically with the Ethovision XT software (Noldus, Wageningen, the Netherlands) or AnyMAZE and the time spent in the two chambers was recorded for the pre- and post-TEST sessions. Subsequently, the preference index was calculated as: time spent in the non-familiar conspecific-paired chamber divided by the time spent in the non-social chamber. Moreover, we calculated the learning index by dividing the preference index calculated at post-TEST by the preference index calculated at pre-TEST.

### 2.3 Drugs

Raclopride (Tocris Bioscience, Tocris House, Bristol, UK), a dopamine D2R/D3R-like receptor antagonist (Köhler et al., 1985), was dissolved in 0.9% saline (vehicle) at conditioning day 1 and stored at −20°C. Raclopride was injected intraperitoneally (i.p.) 20 minutes before the conditioning in a volume of 160 *µl* at a dose of 1.2 mg/kg. Doses and pretreatment interval were chosen based on previous studies (Pina and Cunningham, 2014; Dickinson et al., 2003).

### 2.4 Methodology for pharmacological experiment

During the conditioning days, experimental animals received either saline i.p. injections (vehicle control group, N = 10; 160 *µ*l) or raclopride (N = 10; 160 *µ*l, 1.2 mg/kg) in their home-cage 20 minutes before the conditioning sessions. On the other hand, 20 minutes before pre- and post-TEST all mice were i.p. injected with saline (**Figure 4A**). All experimental animals were housed in groups of 2-3 per cage.

### 2.5 Stereotaxic injections

Purified scrShank3 and shShank3 (AAV1-GFP-U6-scrmb-shRNA; titer: 5.9 × 10^13^ GC/mL and AAV5-ZacF-U6-luc-zsGreen-shShank3; titer: 7.4 × 10^13^ GC/mL, VectorBioLab) injections were delivered in mice younger than P6 as previously described (Bariselli et al., 2016b). After anesthesia induction with a mixture of isoflurane/O_2_, C57Bl/6j wildtype pups were placed on a stereotaxic frame (Angle One; Leica, Germany) and a single craniotomy was made over the VTA. In order to obtain bilateral VTA infection, 200 nl of viral solution was injected at the following coordinates: ML: 0.15 mm, AP: 0.1 mm, DV: −4.0 and −3.9 mm from lambda through a graduated glass pipette (Drummond Scientific Company, Broomall, PA). After behavioral experiments, post hoc analysis was performed to validate the localization of the infection (**Figure 6B**).

### 2.6 Statistical analysis

Analysis of preference index, learning index and time in each chamber was conducted by performing Shapiro-Wilk analysis to assess the normality of sample distributions. T-test with Welch’s correction and paired t-test were used for comparisons between two sample groups when appropriate. When the normality was violated non-parametric Mann-Whitney and Wilcoxon rank-tests were applied. For multifactorial analysis, repeated-measures two-way ANOVA was performed, and P values of main effects and interaction were reported in the figure legends for each experiment. After significant main effects and interactions were revealed, Bonferroni post hoc tests were used for between/within group comparisons and reported in the figure legends or graphs. Significance level was set at P < 0.05. Statistical outliers were excluded when the time spent in either chamber during apparatus exploration (at pre-TEST, post-TEST or extinction sessions) was deviating for more than 2 standard deviations from the group mean. Within this manuscript, the non-familiar conspecific-induced CPP protocol was replicated 4 times in 4 different and independent animal batches.

## 3 Results

### 3.1 Non-familiar conspecific-induced CPP assesses the reinforcing properties of social interaction

To study the reinforcing properties of social interaction, we modified previously published social-induced CPP protocols to obtain CPP mediated by interaction with a non-familiar conspecific in late adolescence male mice. The paradigm started with a pre-TEST phase, during which mice explored for 15 minutes an empty apparatus consisting of two chambers characterized by different contextual cues (**Figure 1A**; see Material and Methods for further details). We then performed 4 days of conditioning, during which mice learnt to associate one compartment of the apparatus with the presence of a non-familiar (novel) social stimulus while the other compartment was left empty (non-social). During each conditioning day, we rapidly alternated for 3 times the exposure to the two chambers (**Figure 1C**). Twenty-four hours after the last conditioning session, we measured the acquisition of a place preference for one of the two chambers by quantifying the time spent exploring each compartment of the empty apparatus (**Figure 1B**). In addition, to compare the reinforcing properties of non-familiar *vs* familiar conspecific, during the conditioning phase, experimental animals encountered in one chamber a non-familiar mouse, while in the other they were exposed to their familiar cage-mate (**Figure 1D**).

### 3.2 The acquisition of non-familiar conspecific-induced CPP does not require single-housing

To determine whether housing conditions affected non-familiar conspecific-induced CPP acquisition, wildtype late adolescent mice underwent conditioning (**Figure 2A**). By measuring the time spent in the non-familiar conspecific-paired chamber and the non-social chamber, we calculated a preference index (time spent in non-familiar conspecific-associated chamber/time spent in the non-social chamber) at the pre- and the post-TEST. We found that single-housed mice increased their preference index at the post-TEST compared to the pre-TEST (**Figure 2B-D**), indicating that they developed a preference for the non-familiar conspecific-associated chamber during conditioning. Additionally, at the post-TEST, single-housed mice spent a significantly longer time in the non-familiar conspecific-paired chamber compared to the non-social one (**Figure 2E**). This indicates that experimental animals were discriminating between contextual cues and expressing a preference for the ones associated with the interaction with a novel conspecific.

**Figure 2.**
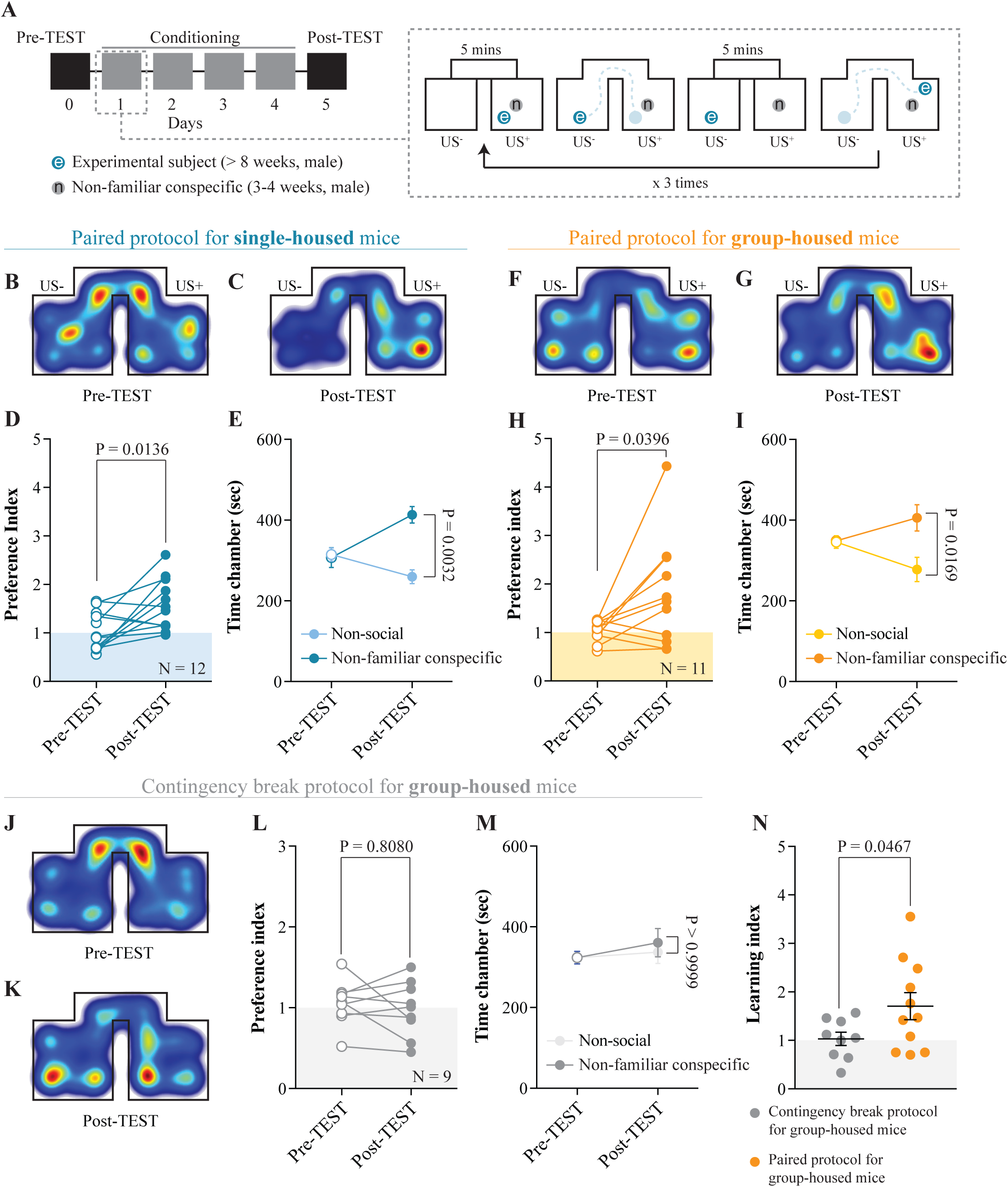
Acquisition of non-familiar conspecific-induced CPP does not require single-housing. (**A**) Left: schematic representation of CPP time-course. Right: schematic representation of a single CPP conditioning alternation. (**B**) Representative apparatus occupancy heat-maps for single-housed mice during pre-TEST and (**C**) post-TEST. (**D**) Preference index calculated at pre- and post-TEST for single-housed mice (N = 12; paired *t*-test: *t*_(11)_ = 2.934; P = 0.0136). (**E**) Time spent in non-social and non-familiar conspecific-paired chambers for single-housed mice during pre- and post-TEST (N = 12; two-way RM ANOVA: time × chamber interaction: *F*_(1, 11)_ = 4.021, P = 0.0702; main effect chamber: *F*_(1, 11)_ = 9.85, P = 0.0094; main effect time: *F*_(1, 11)_ = 6.363, P = 0.0284; followed by Bonferroni’s multiple comparisons test). (**F**) Representative apparatus occupancy heat-maps for group-housed animals at pre-TEST and (**G**) post-TEST. (**H**) Preference index calculated at pre- and post-TEST for group-housed mice (N = 11; paired *t*-test: *t*_(10)_ = 2.366; P = 0.0396). (**I**) Time spent in the non-social and non-familiar conspecific associated chambers for group-housed mice during pre- and post-TEST (N = 11; two-way RM ANOVA: time × chamber interaction: *F*_(1, 10)_ = 5.115, P = 0.0472; main effect chamber: *F*_(1, 11)_ = 3.019, P = 0.1129; main effect time: *F*_(1, 10)_ = 0.2936, P = 0.5998; followed by Bonferroni’s multiple comparisons test). (**J**) Representative apparatus occupancy heat-maps of group-housed mice subject to CPP contingency break at pre-TEST and (**K**) post-TEST. (**L**) Preference index of group-housed mice subjected to CPP contingency break calculated at pre- and post-TEST (N = 9; paired *t*-test: *t*_(8)_ = 0.6537; P = 0.5316). (**M**) Time spent in the two chambers of the apparatus for group-housed mice subjected to CPP contingency break during pre- and post-TEST (the chambers were assigned a non-social and non-familiar property for analyses purposes) (N = 9; two-way RM ANOVA: time × chamber interaction: *F*_(1, 8)_ = 0.1864, P = 0.6773; main effect chamber: *F*_(1, 8)_ = 0.1024, P = 0.7572; main effect time: *F*_(1, 8)_ = 2.401, P = 0.1599; followed by Bonferroni’s multiple comparisons test). (**N**) Learning index calculated for group-housed mice subjected to either a contingency break (N = 9) or a paired protocol CPP (N = 11; unpaired *t*-test with Welch’s correction: *t*_(14,4)_ = 2.175; P = 0.0467). N indicates number of mice. Abbreviations: US = unconditioned stimulus.

Subsequently, we subjected group-housed mice to the same conditioning paradigm used for single-housed animals. Surprisingly, we found that group-housed mice increased their preference for non-familiar conspecific-associated chamber (**Figure 2F-H**) and discriminated between non-familiar conspecific-associated *vs* non-social chamber at the post-TEST (**Figure 2I**). Thus, the non-familiar conspecific-induced CPP paradigm, which relies on the direct and free exploration of a younger and non-familiar conspecific stimulus, allows us to study the reinforcing properties of social interaction without the need to single-house the experimental animals.

To prove the necessity of a contingency between contextual cues and social exposure for the acquisition of non-familiar conspecific-induced CPP, group-housed mice were exposed to the same number of social stimuli presentations, but in alternating chambers (contingency break) during the conditioning phase. In this case, we assigned the non-familiar conspecific-paired chamber as the first chamber in which they were exposed to the non-familiar social stimulus. Under these circumstances, the experimental animals did not increase their preference index (**Figure 2J-L**) and did not show any preference for one of the two chambers at the post-TEST (**Figure 2M**). Finally, to quantify the increase in place preference between pre- and post-TEST and to allow between-group comparisons, we calculated a learning index (preference index at the post-TEST/preference index at the pre-TEST). Confirming the validity of the non-familiar conspecific-induced CPP protocol, group-housed mice conditioned with pairing protocol displayed a significantly higher learning index compared to animals subjected to the contingency break protocol (**Figure 2N**).

### 3.3 Non-familiar conspecific interaction is *per se* reinforcing

To understand whether the novelty component associated to non-familiar social stimuli is necessary to promote contextual learning, we assessed whether group-housed mice would develop any preference for either a familiar or a non-familiar mouse-paired context. By subjecting the animals to conditioning sessions in which they interacted with a non-familiar and a familiar stimulus in different chambers (**Figure 1D**, **3A**), we observed an increase in the preference for the non-familiar conspecific-associated one (**Figure 3B-D**). Additionally, at the post-TEST, mice spent more time in the non-familiar conspecific-paired compartment compared to the one paired with a familiar mouse (**Figure 3E**). Finally, by comparing the learning index obtained from pairing protocols with either non-social session or familiar session, we found no differences in place preference acquisition (**Figure 3F**). Altogether, these results indicate that the social interaction with a non-familiar conspecific is *per se* reinforcing and that CPP is neither driven by social familiarity or non-social pairing.

**Figure 3.**
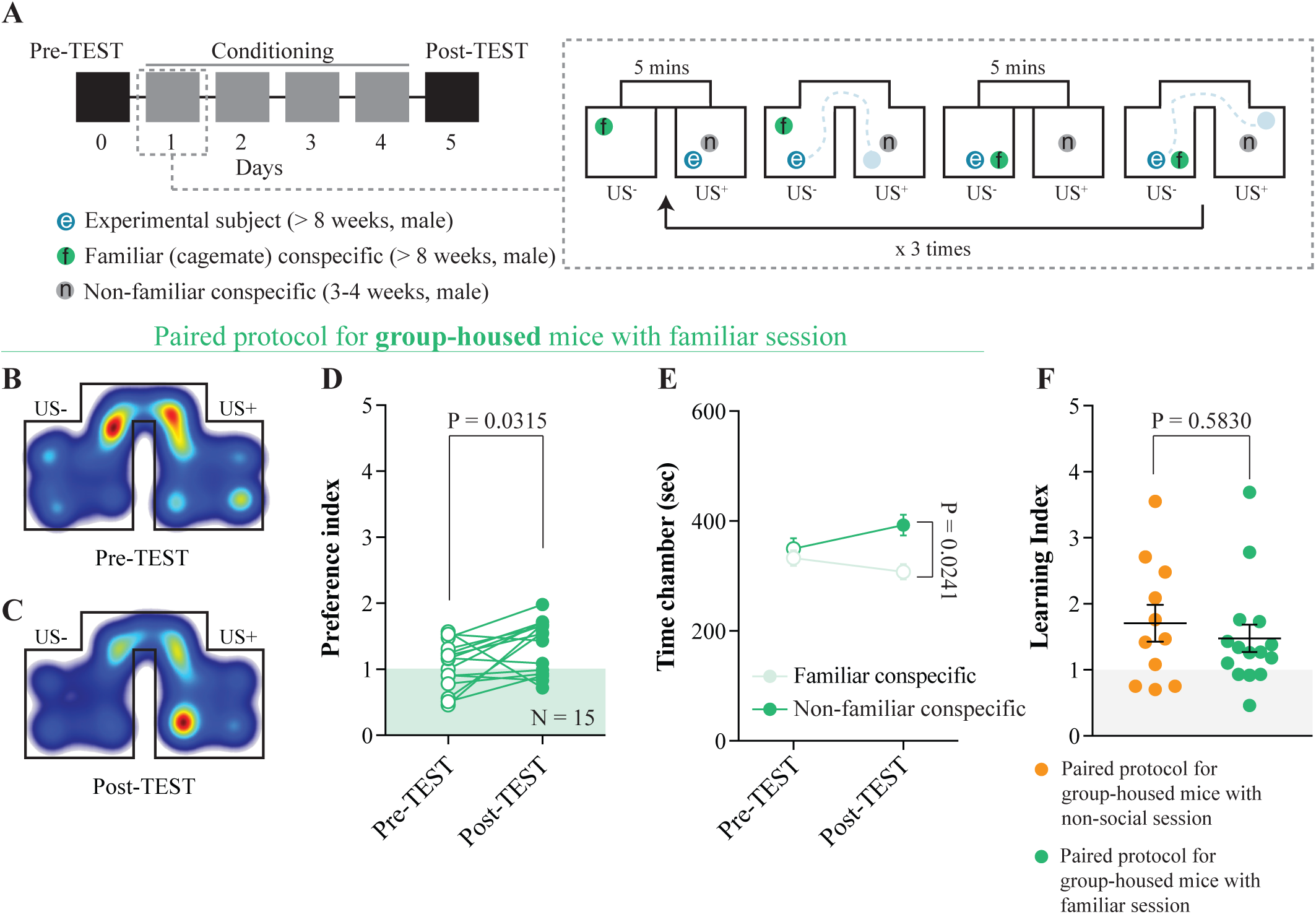
Non-familiar conspecific interaction is *per se* reinforcing. (**A**) Left: schematic representation of CPP time-course. Right: schematic representation of a single CPP conditioning alternation between a familiar session and a non-familiar conspecific session. (**B**) Representative apparatus occupancy heat-maps during pre-TEST and (**C**) post-TEST for group-housed mice subjected to this protocol. (**D**) Preference index calculated at pre- and post-TEST for group-housed mice subjected to this protocol (N = 15; paired *t*-test: *t*_(14)_ = 2.389; P = 0.0315). (**E**) Time spent in familiar and non-familiar conspecific-paired chambers for group-housed mice subjected to this protocol during pre- and post-TEST (N = 15; two-way RM ANOVA: time × chamber interaction: *F*_(1, 14)_ = 4.525, P = 0.0517; main effect chamber: *F*_(1, 14)_ = 3.416, P = 0.0858; main effect time: *F*_(1, 14)_ = 1.657, P = 0.2189; followed by Bonferroni’s multiple comparisons test). (**F**) Learning index calculated for group-housed mice subjected to either the protocol reported in figure 2A (N = 11) or the protocol reported here in figure 3A (N = 15; Mann Whitney test; P = 0.5830). N indicates number of mice. Abbreviations: US = unconditioned stimulus.

### 3.4 D2/D3R function is essential for the acquisition of non-familiar conspecific-induced CPP

Considering the involvement of D2R in controlling both reward-induced place preference acquisition and the expression of appropriate social behavior, we aimed to determine its role in the formation of contextual associations induced by non-familiar conspecific interaction. Group-housed male mice underwent non-familiar conspecific-induced CPP as previously described with the exception that, in this case, they were divided into two groups: after the pre-TEST, one group received i.p. injections of the D2/D3R antagonist raclopride (1.2 mg/kg) while the other received vehicle (saline) treatment, 20 minutes before each conditioning session (**Figure 4A**). By replicating our previous results in a new cohort of mice, we found that vehicle-treated animals increased their preference index (**Figure 4B-D**). Additionally, at post-TEST they explored for longer the non-familiar conspecific-paired chamber compared to the non-social one (**Figure 4E**). However, the raclopride-treated mice failed to increase their preference index for non-familiar social stimulus associated chamber (**Figure 4F-H**), and at post-TEST, had no preference for the non-familiar conspecific-paired chamber or the non-social one (**Figure 4I**). Finally, learning index was significantly reduced by raclopride compared to vehicle treatment (**Figure 4J**). Altogether, these results demonstrate that D2/D3R activation is necessary for the acquisition of non-familiar conspecific-induced CPP.

**Figure 4.**
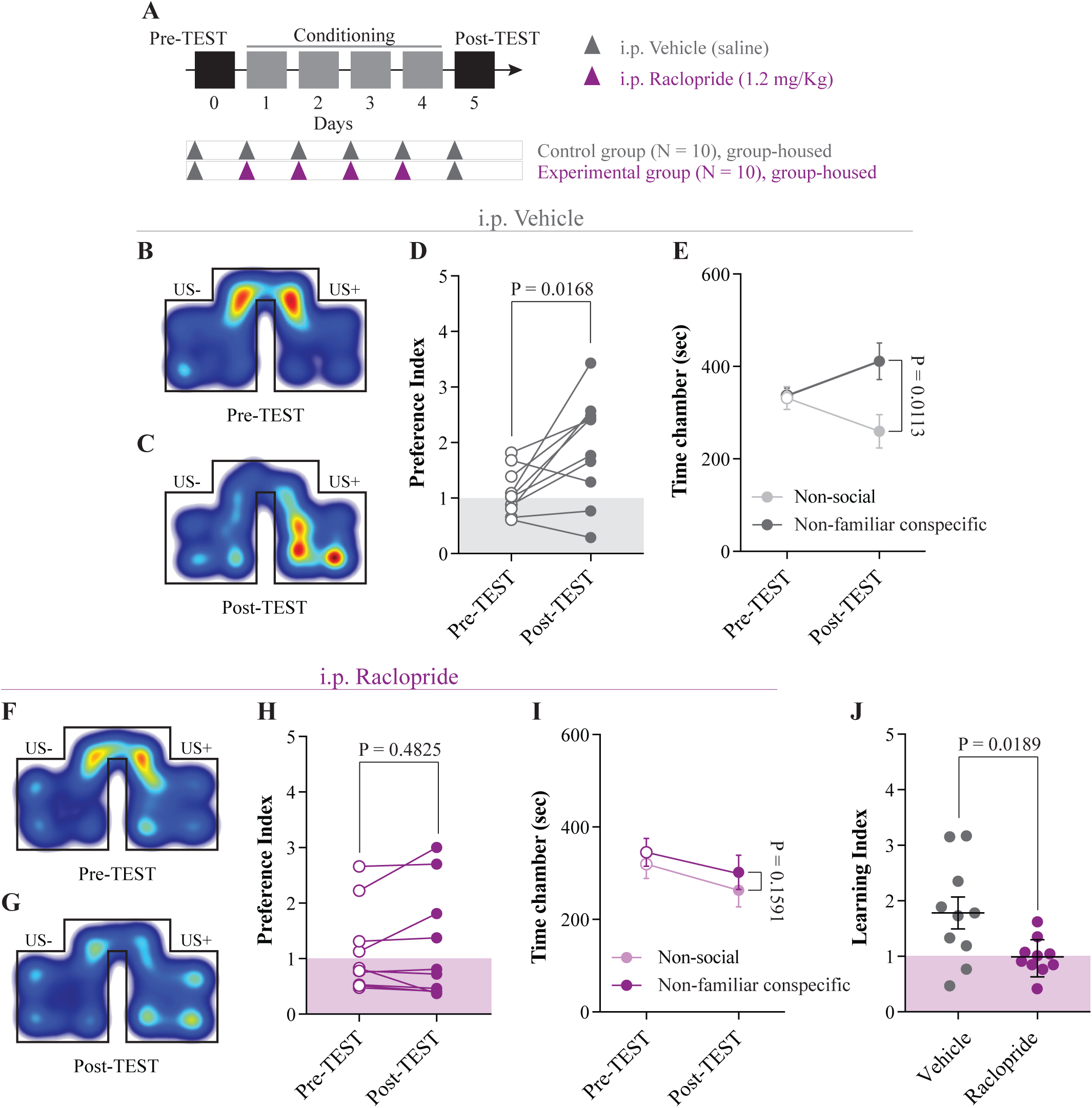
D2R/D3R function is essential for the acquisition of non-familiar conspecific-induced CPP. (**A**) Schematic representation of CPP paradigm and i.p. injections of raclopride and vehicle during conditioning. (**B**) Representative occupancy heat-maps for vehicle-treated group-housed mice during pre-TEST and (**C**) post-TEST. (**D**) Preference index for vehicle-treated group-housed mice calculated at pre- and post-TEST (N = 10; paired *t*-test: *t*_(9)_ = 2.93; P = 0.0168). (**E**) Time spent in non-social and non-familiar conspecific-paired chambers during pre- and post-TEST for vehicle-treated group-housed mice (N = 10; two-way RM ANOVA: time × chamber interaction: *F*_(1, 9)_ = 6.061, P = 0.0360; main effect chamber: *F*_(1, 9)_ = 2.413, P = 0.1548; main effect time: *F*_(1, 9)_ = 0.0146, P = 0.09065; followed by Bonferroni’s multiple comparisons test). (**F**) Representative occupancy heat-maps for raclopride-treated group-housed mice at pre- and (**G**) post-TEST. (**H**) Preference index calculated at pre- and post-TEST for raclopride-treated group-housed mice (N = 10; paired *t*-test: t_(9)_ = 0.7325; P = 0.4825). (**I**) Time spent in non-social and non-familiar conspecific-paired chambers for raclopride-treated group-housed mice during pre- and post-TEST (N = 10; two-way RM ANOVA: time × chamber interaction: *F*_(1, 9)_ = 0.1045, P = 0.7539; main effect chamber: *F*_(1, 9)_ = 0.3304, P = 5795; main effect time: *F*_(1, 9)_ = 19.66, P = 0.0016; followed by Bonferroni’s multiple comparisons test). (J) Learning index of group-housed mice treated with vehicle (N = 10) or raclopride (N = 10) during conditioning phase (unpaired *t*-test: *t*_(18)_ = 2.581; P = 0.0189). N indicates number of mice. Abbreviations: US = unconditioned stimulus.

### 3.5 Extinction of non-familiar conspecific-induced CPP is affected by housing condition

Considering that, after the post-TEST, in adolescent rats social-induced CPP responses are lost within 3 single exposures to the empty CPP apparatus (Trezza et al., 2009), we investigated whether the non-familiar conspecific-induced conditioned responses can also extinguish. Thus, we exposed our experimental animals to 6 extinction sessions, as previously reported (Trezza et al., 2009). Starting 48 hours after the post-TEST, both single-housed and group-housed mice freely explored the empty apparatus for 15 minutes once per day over 6 days (**Figure 5A**). While place preference in group-housed mice was lost after 6 extinction sessions (**Figure 5B**), single-housed mice discriminated between non-familiar and non-social chamber and maintained their preference for the non-familiar conspecific-paired compartment up to extinction session 6 (**Figure 5C, D**). These results indicate that while housing conditions do not induce major changes in the acquisition of the non-familiar conspecific-induced CPP, they affect the extinction of contextual associations induced by free interaction with a novel conspecific.

**Figure 5.**
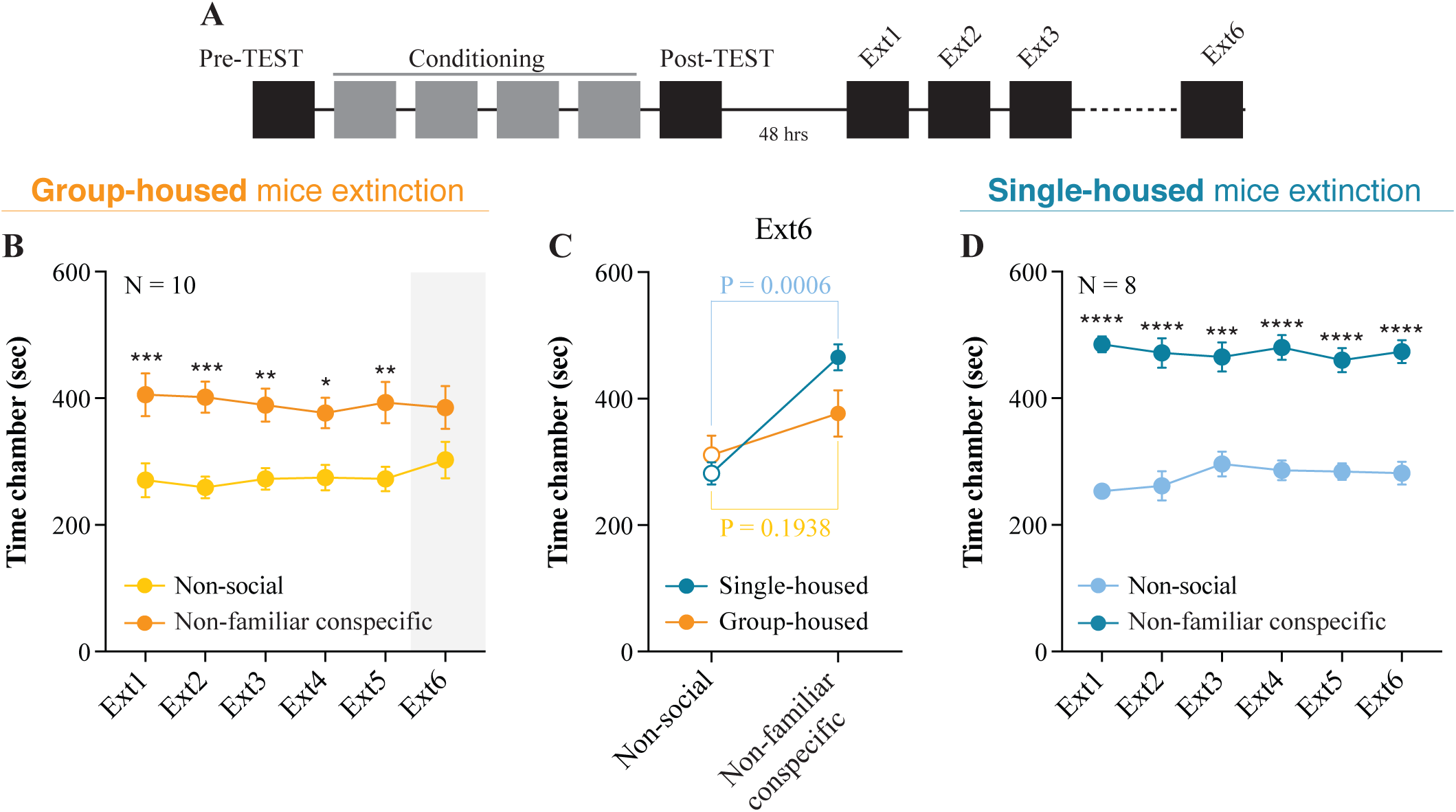
Extinction of non-familiar conspecific-induced CPP is affected by housing condition. (**A**) Schematic representation of CPP acquisition and extinction. (**B**) Time spent in non-social and non-familiar conspecific-paired chambers across extinction sessions for group-housed mice subjected to CPP (N = 10; two-way RM ANOVA: time × chamber interaction: *F*_(5, 45)_ = 0.4486, P = 0.8120; main effect chamber: *F*_(1, 9)_ = 8.484, P = 0.0172; main effect time: *F*_(5, 45)_ = 1.25, P = 0.3021; followed by Bonferroni’s multiple comparisons test). (**C**) Time spent in the non-social or non-familiar conspecific-paired chamber at extinction session 6 for single-housed (N = 8) and group-housed (N = 10) mice (two-way ANOVA: time × group: *F*_(1, 32)_ = 3.316, P = 0.0780; main effect group: *F*_(1, 32)_ = 1.149, P = 0.2917; main effect time: *F*_(1, 32)_ = 23.09, P < 0.0001; followed by Bonferroni’s multiple comparisons test). (**D**) Time spent in non-social and non-familiar conspecific-paired chambers across extinction sessions for single-housed mice (N = 8) subjected to a paired protocol (two-way RM ANOVA: time × chamber interaction: *F*_(5, 35)_ = 1.037, P = 0.4113; main effect chamber: *F*_(1, 7)_ = 78.38, P < 0.0001; main effect time: *F*_(5, 35)_ = 0.997, P = 0.4339; followed by Bonferroni’s multiple comparisons test). N indicate number of mice. Abbreviations: * = p < 0.05, ** = p < 0.01, *** = p < 0.001, **** = p < 0.0001.

### 3.6 VTA SHANK3 downregulation alters the extinction of non-familiar conspecific-induced CPP

Post-natal downregulation of the ASD-related protein SHANK3 in the VTA induces deficits in synaptic transmission, plasticity and social preference dynamics (Bariselli et al., 2016). Since both the excitatory transmission and plasticity at those inputs are essential for associative learning and extinction of reward-induced conditioned responses, we investigated the effects of VTA *Shank3* downregulation on the acquisition and extinction of non-familiar conspecific-induced CPP. More precisely, we downregulated *Shank3* in the VTA of neonatal mice (as previously described in Bariselli et al., 2016; **Figure 6A),** and we group-housed them after weaning for about 5 weeks. At 8 weeks of age (late adolescence), VTA-scrShank3 and VTA-shShank3 mice underwent non-familiar conspecific CPP acquisition and extinction and were subsequently sacrificed for histological validation (**Figure 6B**). We found that VTA-scrShank3 mice expressed a preference for non-familiar conspecific-paired chamber (**Figure 6C-E**) and displayed higher exploration time for non-familiar conspecific-associated chamber at post-TEST (**Figure 6F**). After 48 hours, VTA-scrShank3 mice underwent extinction protocol and, by the extinction session 5, they no longer displayed preference for the non-familiar conspecific-paired chamber (**Figure 6F**). When VTA-shShank3 mice were conditioned, they showed a trend towards a higher preference index between pre- and post-TEST (**Figure 6G-I**), and a significant preference for the non-familiar conspecific-associated chamber *vs* the nonsocial chamber at the post-TEST (**Figure 6J**). Since the learning index of VTA-shShank3 mice was comparable to the one measured from VTA-scrShank3, we concluded that *Shank3* downregulation in the VTA was not inducing major abnormalities in the acquisition of non-familiar conspecific-induced CPP (**Figure 6K**). However, when VTA-shShank3 mice underwent extinction, we noticed that their preference for the non-familiar conspecific-paired chamber was not stable over time and already not significant by the fourth extinction session (**Figure 6J**). To directly compare the extinction of non-familiar conspecific-induced place preference between VTA-scrShank3 and VTA-shShank3 mice, we compared the preference index across pre-TEST, post-TEST and extinction sessions (**Figure 6L**). We found that VTA-shShank3, compared to VTA-scrShank3, had a significantly lower preference index at early extinction sessions indicating an accelerated extinction (**Figure 6L**).

**Figure 6.**
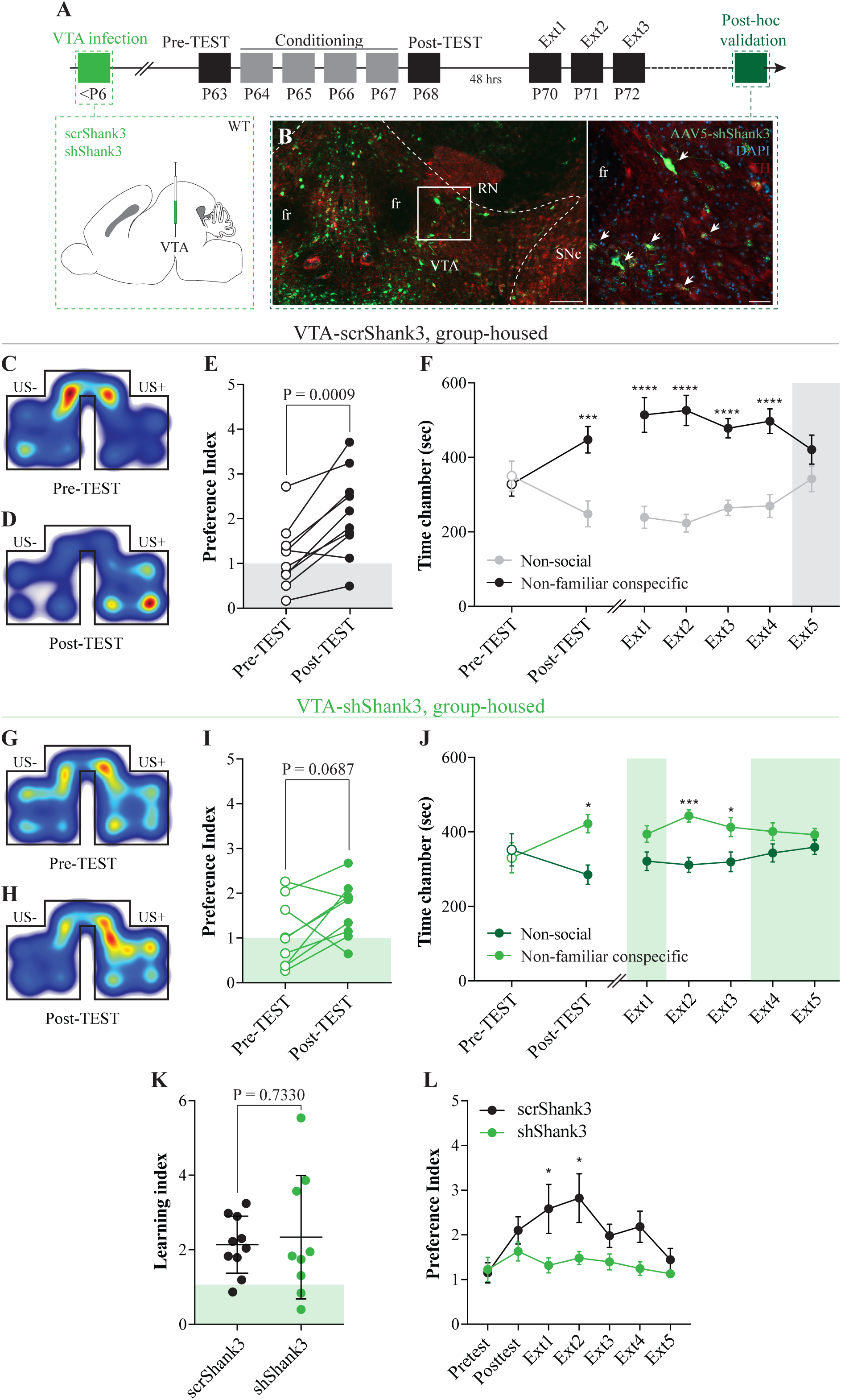
Post-natal VTA *Shank3* downregulation alters extinction of non-familiar conspecific-induced CPP. (**A**) Experimental time course of AAV-shShank3/AAVscrShank3 VTA infection, CPP acquisition, extinction and *post hoc* histological verification of viral infection. **(B)** Histological validation of VTA infection (scale bar: left panel: 150 *µ*m, right panel: 50 *µ*m white arrows indicate TH-shShank3 colocalization). **(C)** Representative occupancy heat-maps for group-housed VTA-scrShank3 mice at pre-TEST and **(D)** post-TEST. **(E)** Preference index calculated at pre- and post-TEST for group-housed VTA-scrShank3 mice (N = 10; paired *t*-test: *t*_(9)_ = 4.827; P = 0.0009). **(F)** Left: time spent in non-social and non-familiar conspecific-paired chamber for group-housed VTA-scrShank3 mice during pre- and post-TEST (N = 10; two-way RM ANOVA: time × chamber interaction: *F*_(1, 9)_ = 25.35, P = 0.0007; main effect chamber: *F*_(1, 9)_ = 1.918, P = 0.1994; main effect time: *F*_(1, 9)_ = 0.4628, P = 0.5134; followed by Bonferroni’s multiple comparisons test). Right: time spent in non-social and non-familiar conspecific-paired chamber for group-housed VTA-scrShank3 mice across extinction sessions (N = 10; two-way RM ANOVA: time × chamber interaction: *F*_(4, 36)_ = 4.624, P = 0.0041; main effect chamber: *F*_(1, 9)_ = 18.32, P = 0.0020; main effect time: *F*_(4, 36)_ = 1.126, P = 0.3595; followed by Bonferroni’s multiple comparisons test). **(G)** Representative occupancy heat-maps for group-housed VTA-shShank3 mice at pre-TEST and **(H)** post-TEST. **(I)** Preference index calculated at pre- and post-TEST for group-housed VTA-shShank3 mice (N = 9; paired *t*-test: *t*_(8)_ = 2.103; P = 0.0687). **(J)** Left: time spent in non-social and non-familiar conspecific-paired chamber for group-housed VTA-shShank3 mice during pre- and post-TEST (N = 9; two-way RM ANOVA: time × chamber interaction: *F*_(1, 8)_ = 4.943, P = 0.0329; main effect chamber: *F*_(1, 8)_ = 2.651, P = 0.1027; main effect time: *F*_(1, 8)_ = 0.119, P = 0.7322; followed by Bonferroni’s multiple comparisons test). Right: time spent in non-social and non-familiar conspecific-paired chambers for group-housed VTA-shShank3 mice during extinction (N = 9; two-way ANOVA: time × chamber interaction: *F*_(4, 32)_ = 1.453, P = 0.2394; main effect chamber: *F*_(1, 8)_ = 6.002, P = 0.0399; main effect time: *F*_(4, 32)_ = 4.226, P = 0.0074; followed by Bonferroni’s multiple comparisons test). **(K)** Learning index calculated for VTA-scrShank3 (N = 10) and VTA-shShank3 (N = 9) (unpaired *t*-test: *t*_(17)_ = 0.3468; P = 0.7330). **(L)** Preference index calculated during pre-TEST, post-TEST and extinction sessions for group-housed VTA-scrShank3 mice (N = 10) and VTA-shShank3 (N = 9) mice (two-way ANOVA: time × group interaction: *F*_(6, 120)_ = 1.439, P = 0.2052; group main effect: *F*_(1, 120)_ = 17.8, P < 0.0001; main effect time: *F*_(6, 120)_ = 2.616, P = 0.0203; followed by Bonferroni’s multiple comparisons test). N indicates number of mice. Abbreviations: US = unconditioned stimulus, fr = fasciculus retroflexus, SNc = substantia nigra pars compacta, RN = red nucleus, * = p < 0.05, ** = p < 0.01, *** = p < 0.001, **** = p < 0.0001.

## 4 Discussion

Reinforcing properties of social interactions have been previously tested by using protocols that require single-housing before and/or during conditioning. Here, by characterizing a modified version of previously published CPP paradigms, we show that while housing conditions (single-housing *vs* group-housing) do not affect the acquisition of a preference for the compartment associated with a non-familiar conspecific, it influences the extinction of non-familiar conspecific-induced contextual associations. Previously, it has been reported that genetic background and neuronal modulators affect acquisition of social-induced CPP (Panksepp et al., 2007; Pinheiro et al., 2016; Dölen et al., 2013). Here, for the first time, we show that acquisition of non-familiar conspecific-induced CPP depends on D2/D3R activation and that post-natal downregulation of *Shank3* in the VTA accelerates its extinction.

### 4.1 D2/D3Rs function is necessary for non-familiar conspecific-induced learning

Within the reward system, midbrain VTA/SNr circuits not only process information related to stimulus novelty in humans (Bunzeck and Duzel, 2006) and control social familiarity perception in mice (Molas et al., 2017), but they also play a major role in associative learning in both non-social and social contexts (Tsai et al., 2009; Adamantidis et al., 2011; Stuber et al., 2008; Hung et al., 2017). Pharmacological blockade of either DA re-uptake or D2R interfere with social-induced CPP in rats (Trezza et al., 2009) and testosterone-induced CPP in male Syrian hamsters (Bell and Sisk, 2013), respectively. In the present study, blocking D2/D3Rs prevented associative learning induced by non-familiar conspecific interaction in late adolescence C57Bl6/j mice, thus identifying a necessary molecular target of DA for the acquisition of non-familiar conspecific-induced CPP. This evidence links the possibility that the genetic variants of the *DRD2/DRD3* and dopamine transporter alleles found in ASD patients (Comings et al., 1991; Staal et al., 2012; Cartier et al., 2015) might be causal to sociability defects reported in these individuals, perhaps due to aberrant social-induced learning. Future studies are needed to investigate the VTA downstream circuits involved in the formation of contextual associations induced by social interaction with a non-familiar conspecific.

### 4.2 Effects of housing on the extinction of contextual associations induced by interaction with a non-familiar conspecific

Social-induced CPP can be lost in adolescent rats (Trezza et al., 2009), and we demonstrated that social-induced conditioned responses could be induced and also lost in late adolescent mice. However, we found that housing conditions affect the extinction of the preference for the non-familiar conspecific-paired chamber. In single-housed compared to group-housed animals, the decreased rate of extinction of these conditioned responses might be due to the increased strength of the association between contextual cues (CS) and the presence of a non-familiar conspecific, which would strongly counteract the new association (CS-absence of social stimuli). The increased strength of the associations in single-housed mice could stem from the increased subjective value of social stimuli during the acquisition of the non-familiar conspecific-induced CPP as a consequence of the absence of social contacts in their home-cage. In fact, the saliency of social events is modulated by social context, as, compared to non-lonely individuals, lonely individuals convey higher attention and accuracy in decoding social cues (Pickett et al., 2004) and have a greater ability to recall socially-related information (Gardner et al., 2005). Alternatively, the absence of social contacts in the home-cage during extinction might favor, in the presence of the contextual cues previously associated with social interaction availability, a behavioral strategy that maximizes the probability of engaging in social interactions (*e.g.* spending more time in the chamber paired with the exposure to a non-familiar conspecific). Also, in this case, the influence of the primary association on the behavioral outcome would be increased at the expense of the new CS-absence of social interaction association, thus resulting in a reduced extinction of the conditioned response. Considering the effects of single-housing on the extinction of non-familiar conspecific-induced CPP, investigating the behavioral and neuronal mal-adaptations induced by social isolation on the acquisition and extinction of social-induced conditioned responses might be beneficial for individuals affected by “loneliness” (*Cacioppo JT & Cacioppo S, The Lancet, “The growing problem of loneliness*”).

### 4.3 *Shank3* downregulation in VTA accelerates extinction of non-familiar conspecific-induced CPP

SHANK3 is a post-synaptic protein that orchestrates excitatory synaptic function and whose mutations are associated with ASD (Jiang and Ehlers, 2013). In recent years, several genetic mouse models have been developed to study the involvement of each variant in synaptic and ASD-like behavioral impairments (Bariselli and Bellone, 2016). We previously found that developmental downregulation of *Shank3* in the VTA promotes an aberrant insertion of non-canonical α-amino-3-hydroxy-5-methyl-4-isoxazolepropionic acid (AMPA) receptors and increases AMPA/NMDA (N-methyl-D-aspartate receptor) ratio at excitatory inputs onto DA neurons. In parallel, VTA-SHANK3 deficiency impaired social preference (Bariselli et al., 2016b), pointing to a deficit in social motivation and reward processing. Here, we tested whether these deficits might be accompanied by impairments in the formation of association between contextual cues and social interaction with a non-familiar conspecific. We found that, although VTA-shShank3 mice discriminated between non-familiar conspecific-paired and non-social compartment at the post-TEST, they showed only a statistical trend towards an increase in their preference for the non-familiar conspecific-paired compartment before and after conditioning. These results might indicate subtle deficits in acquisition of non-familiar conspecific-induced contextual association. However, when VTA-shShank3 mice were repeatedly exposed to contextual cues in the absence of the social stimulus animal, their preference for non-familiar conspecific-paired chamber became erratic and revealed an accelerated rate of extinction. This effect could be due to a reduced strength of the association as a consequence of *Shank3* downregulation in the VTA. According to the hypothesis that extinction is learning of a new association, it is also plausible that in VTA-shShank3 mice the new association might easily overpower the weak primary association, thus resulting in an accelerated disappearance of the conditioned responses. To our knowledge this is the first time that an extinction deficit is reported in an ASD-related genetic dysfunction restricted to the VTA, but further investigation is warranted to understand the behavioral and neurobiological mechanisms underlying aberrant learning and extinction of those associations.

### Considerations on non-familiar conspecific-induced CPP

We tested the formation of contextual associations induced by social interaction with a non-familiar conspecific in late adolescent male (8/9 weeks old) mice using early adolescence males (3-4 weeks old) as social stimuli to minimize antagonistic behavior. However, considering that offensive behavior is a trait expressed by C57Bl/6 male mice (Crawley et al., 1997) and that aggression promotes place preference in CD1 male mice (Golden et al., 2016), further investigation is needed to address the role of offensive behavior in the acquisition of non-familiar conspecific-induced CPP in C57Bl/6j subjects. Additionally, considering the great impact of sexual hormones on the acquisition of reward-induced place preference (Calipari et al., 2017), future studies will need to assess the acquisition of non-familiar conspecific-induced place preference in female mice at various stages of their estrous cycle.

Social novelty recognition induces territorial urinary marking (Arakawa et al., 2008), increases exploratory behavior (Ferguson et al., 2000), and produces exploratory preference, which is only expressed during the first five minutes of the three-chamber task (Nadler et al., 2004). For these reasons, during the conditioning phase, we exposed the experimental animals to non-familiar social stimuli for only five minutes. This maximizes the number of pairing sessions per day, which is positively associated to the development of social-induced CPP (Thiel et al., 2008; Trezza et al., 2009). We found that under these circumstances, the acquisition of association between contextual cues and interaction with a non-familiar conspecific does not require single-housing. Moreover, non-familiar conspecific-induced CPP is expressed when group-housed mice are conditioned with familiar *vs* non-familiar social stimuli, indicating that social interaction with non-familiar conspecific is *per se* reinforcing. This evidence contrasts with other studies that show single-housing as a necessary condition for social-induced CPP acquisition in rats (Trezza et al., 2009; Douglas et al., 2004), possibly due to differences in experimental conditions and protocol. Thus, further experiments are needed to investigate the modulation of social-induced CPP acquisition by species and age of the experimental subjects, number and duration of conditioning sessions, the sociability of stimuli mice and hierarchy dynamics within the home-cage of both experimental and stimulus subjects.

In conclusion, the CPP is a behavioral procedure classically used to assess the reinforcing properties of drugs of abuse and natural rewards. The observation of social-induced CPP in group-housed mice demonstrates that direct social interaction with a non-familiar conspecific can be reinforcing and can be studied without the confounding behavioral and neurobiological mal-adaptations induced by social isolation. Extinction and reinstatement of drug-induced CPP responses are behavioral procedures used to assess “drug-seeking” behavior in mice (Itzhak and Martin, 2002). For this reason, many studies have focused their attention on the cellular, synaptic and circuit adaptations that underlie “drug-seeking” behavior to facilitate extinction, to reduce reinstatement (Prast et al., 2014; Thanos et al., 2009; Voigt et al., 2011; Malvaez et al., 2013) and, ultimately, to aid the research for treating addiction. Similarly, the methods of extinction, and possibly reinstatement, of non-familiar conspecific-induced CPP responses are a way to observe and assess “social-seeking” behavior in mice. This paradigm could prove instrumental in identifying mechanisms underlying “social-seeking”, and help to find pharmacological targets to promote social-induced learning.

## 5 Conflict of interest

The authors declare that the research was conducted in the absence of any commercial or financial relationships that could be construed as a potential conflict of interest.

## 6 Author contributions

SB, AC and ST performed the behavioural experiments. SB and AC prepared the figures. SB and CB designed the study and wrote the manuscript.

## 7 Funding

C.B is supported by the Swiss National Science Foundation, Pierre Mercier Foundation, and NCCR Synapsy.

## 8 Acknowledgements

We thank Jeremy Pantone, Giulia Casarotto and Lorena Jourdain for their technical contributions to the behavioural experiments and the histological validation.

